# *Artemisia annua* and *Artemisia afra* extracts exhibit strong bactericidal activity against *Mycobacterium tuberculosis*

**DOI:** 10.1101/2020.04.26.062331

**Authors:** Maria Carla Martini, Tianbi Zhang, John T. Williams, Robert B. Abramovitch, Pamela J. Weathers, Scarlet S. Shell

## Abstract

**Ethnopharmacological relevance:** Emergence of drug-resistant and multidrug-resistant *Mycobacterium tuberculosis* (Mtb) strains is a major barrier to tuberculosis (TB) eradication, as it leads to longer treatment regimens and in many cases treatment failure. Thus, there is an urgent need to explore new TB drugs and combinations, in order to shorten TB treatment and improve outcomes. Here, we evaluate the potential of two medicinal plants, *Artemisia annua*, a natural source of artemisinin (AN), and *Artemisia afra*, as sources of novel antitubercular agents.

**Aim of the study:** Our goal was to measure the activity of *A. annua* and *A. afra* extracts against Mtb as potential natural and inexpensive therapies for TB treatment, or as sources of compounds that could be further developed into effective treatments.

**Materials and Methods:** The minimum inhibitory concentrations (MICs) of *A. annua* and *A. afra* dichloromethane extracts were determined, and concentrations above the MICs were used to evaluate their ability to kill Mtb and *Mycobacterium abscessus in vitro*.

**Results:** Previous studies showed that *A. annua* and *A. afra* inhibit Mtb growth. Here, we show for the first time that *Artemisia* extracts have a strong bactericidal activity against Mtb. The killing effect of *A. annua* was much stronger than equivalent concentrations of pure AN, suggesting that *A. annua* extracts kill Mtb through a combination of AN and additional compounds. *A. afra*, which produces very little AN, displayed bactericidal activity against Mtb that was substantial but weaker than that of *A. annua*. In addition, we measured the activity of *Artemisia* extracts against *Mycobacterium abscessus*. Interestingly, we observed that while *A. annua* is not bactericidal, it inhibits growth of *M. abscessus*, highlighting the potential of this plant in combinatory therapies to treat *M. abscessus* infections.

**Conclusion:** Our results indicate that *Artemisia* extracts have an enormous potential for treatment of TB and *M. abscessus* infections, and that these plants contain bactericidal compounds in addition to AN. Combination of extracts with existing antibiotics may not only improve treatment outcomes but also reduce the emergence of resistance to other drugs.

## 1. INTRODUCTION

3,000 years after the first documented case of tuberculosis (TB) (Barberis et al., 2017) and 130 years after the discovery that *Mycobacterium tuberculosis* (Mtb) is the causative agent of TB, this disease remains one of the major worldwide health challenges. In 2018 alone, 10 million people fell ill with TB and 1.2 million died from the disease, positioning TB as one of the top 10 causes of death worldwide (WHO, 2019). A major barrier to lowering this number is the suboptimal nature of TB antibiotic therapies. Drug-sensitive TB must be treated with six months of combination therapy to prevent relapse and minimize the emergence of resistance. Drug-resistant TB requires even longer treatment regimens with more debilitating side effects and poorer outcomes. Thus, better drugs and combinations are needed to make TB treatment faster and less toxic. Since the late 1970s, only four drugs (linezolid, bedaquiline, delamanid and pretomanid) have been made available as second-line antitubercular agents to treat multidrug-resistant and extensively drug-resistant TB (Keam, 2019; Lee et al., 2012; Osborne, 2013; Ryan and Lo, 2014). Despite the recent introduction of these antibiotics in TB treatment, bedaquiline and delamanid resistance have already been reported in Mtb clinical isolates (Mokrousov et al., 2019; Polsfuss et al., 2019), indicating that resistance emerges quickly and highlighting the urgent need to develop new drugs and combinations to improve TB therapy.

Recent work presented the antimalarial drug artemisinin (AN) as a promising antitubercular drug (Choi, 2017; Zheng et al., 2017). AN inhibits Mtb survival of hypoxia *in vitro* by blocking the DosRST two-component regulatory system, necessary for survival of Mtb during non-replicating persistence (Zheng et al., 2017; Zheng et al., 2019). It also has bactericidal activity against Mtb during aerated growth for reasons that are not fully elucidated but may involve lipid peroxidation (Patel et al., 2019). *Artemisia annua* is the natural source of AN, which was the scaffold for development of semi-synthetic derivatives now in widespread clinical use for treatment of malaria. Various *Artemisia* species are used in traditional medicine around the world, including use of *A. afra* in southern Africa to treat fever and cough, classic sympotoms of TB (Thring and Weitz, 2006). This traditional usage has prompted studies testing *Artemisia* extracts for activity against numerous pathogens and conditions, including mycobacteria in culture and in a murine model of tuberculosis (Cantrell et al., 1998; Uba et al., 2003). In addition, *A. afra*, which produces little to no AN, displayed inhibitory activity against Mtb (Mativandlela et al., 2008; Ntutela et al., 2009) suggesting that compounds other than AN in the extract can inhibit Mtb growth.

In the present study, we measured the ability of *A. annua* and *A. afra* extracts to kill Mtb in culture. We demonstrate that both extracts are strongly bactericidal against Mtb and produced more killing that equivalent concentrations of pure AN. We also tested the impact of these extracts on the emerging pathogen *M. abscessus* and the model organism *M. smegmatis*, and found that extracts inhibited growth but were not bactericidal at the concentrations tested.

## 2. MATERIALS AND METHODS

### 2.1 Strains and growth conditions

*M. tuberculosis* mc^2^6230 (Δ*panCD*, ΔRD1, (Sambandamurthy et al., 2006)) and virulent Erdman strains were grown in Middlebrook 7H9 supplemented with OADC (Oleic acid Albumin Dextrose Catalase, final concentrations 5 g/L bovine serum albumin fraction V, 2 g/L dextrose, 0.85 g/L sodium chloride, and 3 mg/L catalase), 0.2% glycerol and 0.05% Tween 80. For the Mtb mc^2^6230 auxotrophic strain, pantothenate was added to a final concentration of 24 μg/mL. Mtb mc^2^6230 and Mtb Erdman were grown in BSL-2 and BSL-3 containments, respectively, in accordance with institutionally approved standard operating procedures established for these strains. *M. smegmatis* mc^2^155 and *M. abscessus* ATCC_19977 strains were grown in Middlebrook 7H9 supplemented with ADC (Albumin Dextrose Catalase, final concentrations 5 g/L bovine serum albumin fraction V, 2 g/L dextrose, 0.85 g/L sodium chloride, and 3 mg/L catalase), 0.2% glycerol and 0.05% Tween 80. *M. abscessus* was grown in BSL-2 containment in accordance with institutionally approved standard operating procedures.

Middlebrook 7H10 OADC solid media supplemented with 0.2% glycerol was used to count colony forming units (CFUs) for all strains. 24 μg/mL pantothenate was added to Mtb mc^2^6230 plates and 10 μg/mL cycloheximide was added to Mtb Erdman plates to prevent fungal contamination.

### 2.2 Preparation of plant extracts

Dried leaves of *A. annua* L. SAM cultivar (voucher MASS 00317314) and *A. afra* Jacq. ex Willd. (SEN) (voucher Université de Liège LG0019529) were used and their phytochemical contents are detailed in (Weathers and Towler, 2014) and (Munyangi et al., 2018), respectively. *A. annua* was propagated in-house and harvested as described (Towler and Weathers, 2015). Dried *A. afra* leaves were obtained from Guy Mergei, Université de Liège, Belgium. Dried leaf powder of *A. annua* and *A. afra* were resuspended in dichloromethane (1 g dried leaves per 20 mL DCM) and extracted under sonication as previously detailed (Desrosiers et al., 2020; Weathers et al., 2014). AN was quantified by GC-MS using the method described in Weathers and Towler (2014) with the following modifications: ion source temperature, 230°C; inlet, 150°C; transfer line, 280°C; oven temperature, 125°C held for 1 min, then increased to 240°C at 5°C/min, and then increased to 300°C at 30°C/min. *A. annua* and *A. afra* extracts used here contained 0.82% and ≤0.026% (w/w) of AN in dry weight, respectively. Dry extracts were sterilized by ethylene oxide, degassed for one day, stored at −20°C, and later resuspended in sterile DMSO for use in experiments.

### 2.3 Determination of minimum inhibitory concentration (MIC)

MICs of AN and *Artemisia* extracts in Mtb strain mc^2^6230 were determined by resazurin microtiter assay (REMA) as previously reported (Choi, 2017) with minor modifications. Briefly, Mtb log-phase cultures were adjusted to a final OD=0.001. Bacterial suspensions were inoculated into 96 well microtiter plate containing final concentrations of i) 1.17-600 μg/mL pure AN or ii) *A. annua* extract containing 1.17-600 μg/mL AN or iii) *A. afra* extract made from equivalent dry weights as the *A. annua* extract. All wells contained 2.5% DMSO and final volumes were 200 μL. Controls consisting of 7H9 medium alone or 7H9 medium + drug/extract or 7H9 medium + bacterial culture were included. Plates were covered with breathable paper and plastic lids, placed in plastic bags and incubated at 37°C and 125 rpm for 7 days. After this time, 20 μL 0.02% (w/v) resazurin solution was added to each well and incubated for 24h. A change in color from blue to pink indicated bacterial growth. The MIC was defined as the lowest concentration of drug/extract that prevented visible color change.

### 2.4 Measurement of plant extract effects on mycobacterial viability

For Mtb mc^2^6230, *M. abscessus*, and *M. smegmatis*, log-phase cultures were sub-cultured to an OD=0.1 and 5 mL aliquots were placed into 50 mL conical tubes. Pure AN, *A. afra*, or *A. annua* extracts were added to achieve the desired concentrations. Cultures containing 2.5% DMSO were included as a control. Cultures were allowed to grow at 37°C and 200 rpm for 14 days (Mtb) or 7 days (*M. abscessus* and *M. smegmatis*). Samples from all treatments were collected at time 0 and at different timepoints and serial dilutions were plated on 7H10 to calculate the number of CFUs. The number of colonies was determined after 40 days (Mtb) or 3 days (*M. abscessus* and *M. smegmatis*) of incubation at 37°C. For Mtb Erdman strain, 30 mL of log-phase cultures were pelleted and resuspended in fresh 7H9 and 5 mL aliquots were placed in T-25 flasks and AN, *A. afra*, or *A. annua* were added. Cultures were incubated at 37°C (+5% CO2) in T-25 flasks without shaking for 12 days. CFUs were determined following the same procedure as with the other strains.

## 3. RESULTS AND DISCUSSION

In order to measure the potential of *Artemisia* extracts to kill Mtb, we first sought to determine the concentrations of pure AN and DCM extracts of *A. annua* and *A. afra* that inhibited growth of Mtb strain mc^2^6230. We found that the MIC for pure AN was 75 μg/mL. For *A. annua* the MIC was the extract from 4.81 mg of dried leaves per mL media, which resulted in 37.5 μg/mL of AN. For *A. afra* the MIC was the extract from 4.81 mg of dried leaves per mL media, which contained <1.3 μg/mL of AN. These results show that *Artemisia* extracts inhibit Mtb growth to an extent that cannot be fully explained by their AN content. The MIC is used to evaluate the antimicrobial efficacy of antibiotics by measuring the bacteriostatic capability of a certain agent, but does not provide information on its bactericidal activity. Previous studies reported growth inhibition by *A. annua* and *A. afra* extracts in Mtb cultures (Cantrell et al., 1998; Mativandlela et al., 2008; Ntutela et al., 2009; Uba et al., 2003). However, the bactericidal activity of these extracts has to our knowledge not yet been reported. To investigate the potential of *A. annua* extract as a bactericidal agent, we treated Mtb mc^2^6230 cultures with concentrations above the MIC of *A. annua* extract and found that while AN alone was bactericidal, the extract produced more killing with faster kinetics than equivalent AN concentrations alone (Fig. 1A). In addition, a two-fold increase in pure AN concentration (150 μg/mL to 300 μg/mL) did not increase killing, while an equivalent increase in *A. annua* concentration remarkably potentiated bactericidal activity against Mtb (Fig 1A). These data suggest that *A. annua* extract kills Mtb through a combination of AN and additional compounds present in the plant extract.

**Figure 1.**
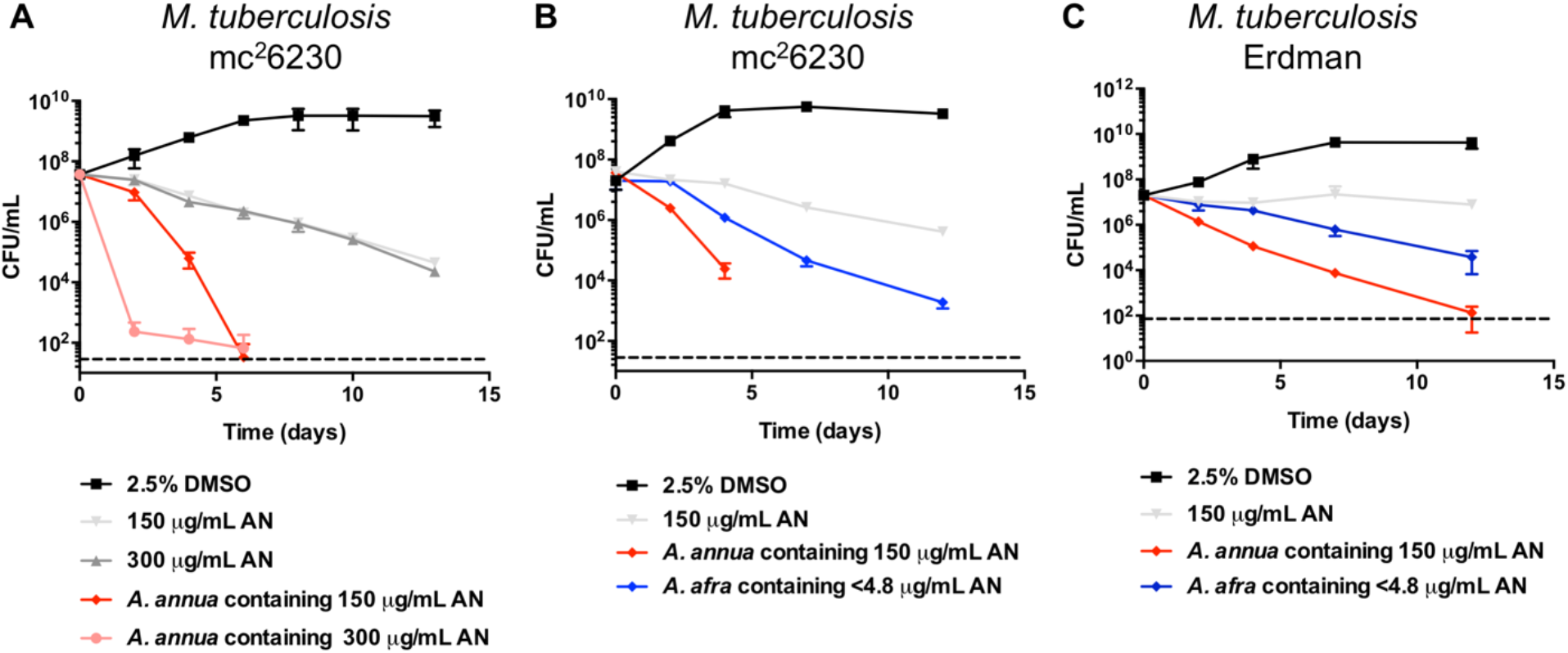
*Artemisia* extracts exhibit strong bactericidal activity against *M. tuberculosis*. **A**. *M. tuberculosis* mc^2^6230 was incubated in the presence of 150 μg/mL or 300 μg/mL of pure AN, or *A. annua* extract containing equivalent concentrations of AN. **B** and **C**. *M. tuberculosis* mc^2^6230 (B) or Erdman (C) was exposed to 150 μg/mL of pure AN or *A. annua* extract containing equivalent concentrations of AN or *A. afra* at equivalent dry weight as the *A. annua* extract. 2.5% DMSO was included as a control.

We further measured the potential of *A. afra* against Mtb mc^2^6230. We found that extracts of this plant exhibited bactericidal activity, although to a lesser extent than extracts of *A. annua* made from an equivalent mass of dried leaves (Fig 1B). Given the much lower levels of AN in *A. afra* compared to *A. annua*, this result suggests that the stronger bactericidal activity of *A. annua* may be due to the combination of AN and other plant compounds. However, we cannot rule out the possibility that the difference is due to differences in other aspects of the phytochemistry of the two species. It is important to highlight that the *A. afra* extract displayed significantly greater killing than pure AN at a concentration >30-fold higher than that present in the extract, which reinforces the premise that other compounds present in *Artemisia* plants contribute to their bactericidal effects.

Similar bactericidal activities of *Artemisia* extracts were observed when the virulent Mtb Erdman strain was used (Fig 1C), although in this case AN prevented Mtb growth but did not display bactericidal activity. The differences in pure AN outcomes as well as the slightly lower killing observed for *Artemisia* extracts in Erdman compared to mc^2^6230 strain may be due to the different experimental conditions used in these assays (see Section 2.4). In addition, mc^2^6230 is a derivative of H37Rv, which has been shown to behave differently than Erdman strain in other aspects (Manabe et al., 2003; North and Izzo, 1993).

We also sought to evaluate the potential of *Artemisia* extracts against *M. abscessus*, a non-tuberculous mycobacterium causing severe infections in immunocompromised patients and whose treatment is very restricted due to the limited number of effective drugs. Interestingly, we found that pure AN and *A. afra* do not hamper *M. abscessus* growth, while *A. annua* showed bacteriostatic activity against this pathogen (Fig 2A). Athough bactericidal activity is highly desirable, there is debate about the extent to which bactericidal drugs are better than bacteriostatic drugs to treat clinical infections (Nemeth et al., 2015; Pankey and Sabath, 2004; Rhee and Gardiner, 2004; Wald-Dickler et al., 2018). Antibiotic efficacy *in vivo* depends on many other factors such as drug combinations, pharmacodynamics, and pharmacokinetics (Rhee and Gardiner, 2004). In addition, some antibiotics have been shown to exhibit bacteriostatic or bactericidal activity, depending on the bacterial growth phase or their interaction with other drugs (Bakker-Woudenberg et al., 2005; Lobritz et al., 2015; Yamori et al., 1992; Zhang et al., 2014). Bacteriostatic antibiotics are effective in treating *M. abscessus* and other mycobacterial infections and their use is also important in preventing emergence of drug resistance (Ferro et al., 2016; Vilchèze and Jacobs, 2012) especially when pharmacological options are limited. Thus, we propose that *A. annua* has potential to treat *M. abscessus* infections and warrants further study.

**Figure 2.**
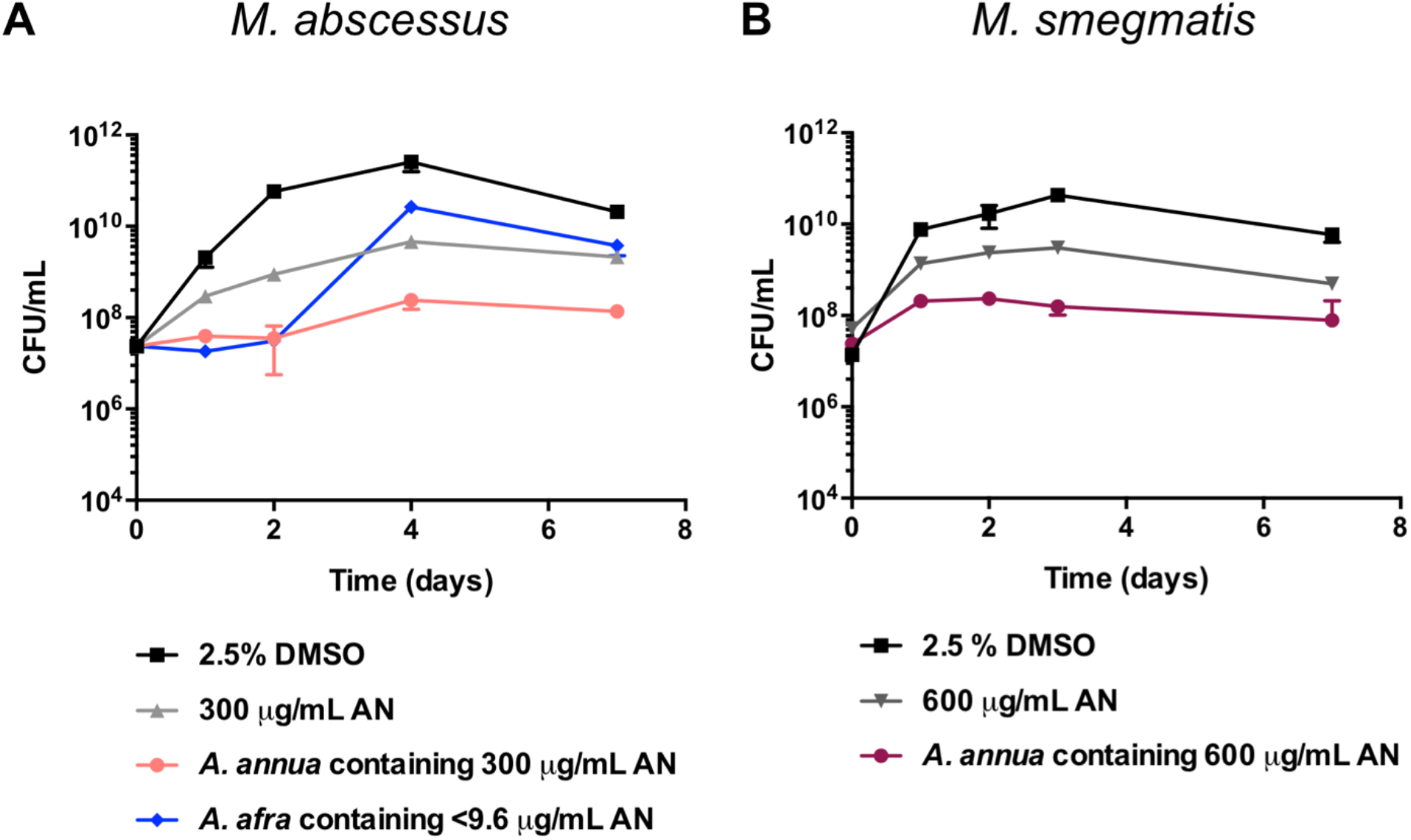
*Artemisia* extracts have different impacts on *M. abscessus* and *M. smegmatis*. The strains were incubated in presence of 300 μg/mL **(A)** or 600 μg/mL **(B)** of pure AN, or *A. annua* extract containing equivalent concentrations of AN, or *A. afra* at equivalent dry weight as the *A. annua* extract. 2.5% DMSO was included as a control.

We finally investigated the effect of *A. annua* extract against *M. smegmatis*, a fast-growing non-pathogenic mycobacterium widely used as a model system to study many aspects of Mtb physiology. We found that, while growth was significantly affected, neither pure AN nor the extract have the ability to fully inhibit growth or kill this organism at the concentrations tested (Fig 2B).

## 4. CONCLUSIONS

The strong bactericidal effect of *A. annua* and *A. afra* extracts against Mtb and the bacteriostatic activity of *A. annua* against *M. abscessus* point out the enormous potential of these extracts, or compounds within them, to treat mycobacterial infections. The stronger killing activity of *A. annua* compared to pure AN at equivalent concentrations and the moderate killing of *A. afra* suggest that other metabolites are important for these bactericidal activities, making these plants an excellent alternative to the use of pure AN. Another aspect to be considered is that using *A. annua* extracts for TB treatment could potentially increase the bioavailability of AN, as we previously observed for malaria treatment in a rat model (Desrosiers et al., 2020). In addition, the implementation of *Artemisia* extracts in Mtb and *M. abscessus* infections treatment could slow down or prevent the emergence of resistance to other drugs. Further study is needed to identify the active phytochemicals in these extracts and evaluate their potential as antitubercular drugs. Additionally, our study focused on plant extracts made with a single solvent. Other solvents should be tested to evaluate the potential of compounds that are not efficiently extracted by DCM.

## AUTHOR CONTRIBUTIONS

M.C.M., S.S.S., P.J.W., and R.B.A. conceived and designed experiments. T.Z. prepared plant extracts. M.C.M., T.Z., and J.T.W. performed antimycobacterial activity assays. M.C.M. and S.S.S. wrote the manuscript.

## ACKNOWLEDGEMENTS

This work was funded in part by R01AI116605 (to RBA) and phytochemical analysis was funded by the National Center for Complementary and Integrative Health, award number NIH-2R15AT008277-02 (to PW). The content is solely the responsibility of the authors and does not necessarily represent the official views of the National Center for Complementary and Integrative Health or the National Institutes of Health. We thank Melissa Towler for assistance with quantification of the artemisinin content in plant extracts. We thank Guy Mergei for providing *A. afra* plant material. We thank members of the Shell and Weathers labs for technical assistance and helpful discussions.

## Abbreviations

Mtb: *Mycobacterium tuberculosis*
TB: tuberculosis
AN: artemisinin
MIC: minimum inhibitory concentration
OD: optical density
CFUs: colony forming units
DCM: dichloromethane.

